# Network diffusion model predicts neurodegeneration in limb-onset amyotrophic lateral sclerosis

**DOI:** 10.1101/2021.07.02.450445

**Authors:** Anjan Bhattarai, Zhaolin Chen, Phyllis Chua, Paul Talman, Susan Mathers, Caron Chapman, James Howe, CM Sarah Lee, Yenni Lie, Govinda R Poudel, Gary F Egan

**Author notes:** **Correspondence to:** Anjan Bhattarai; Monash Biomedical Imaging, Monash University, 770 Blackburn Rd, Clayton, VIC 3168, Australia. Govinda R Poudel and Gary F Egan contributed equally to this work.

## Abstract

The trans-neural propagation of phosphorylated 43-kDa transactive response DNA-binding protein (pTDP-43) contributes to neurodegeneration in Amyotrophic Lateral Sclerosis (ALS). We investigated whether Network Diffusion Model (NDM), a biophysical model of spread of pathology via the brain connectome, could capture the severity and progression of neurodegeneration (atrophy) in ALS. We measured degeneration in limb-onset ALS patients (n=14 at baseline, 12 at 6-months, and 9 at 12 months) and controls (n=12 at baseline) using FreeSurfer analysis on the structural T1-weighted Magnetic Resonance Imaging (MRI) data. The NDM was simulated on the canonical structural connectome from the IIT Human Brain Atlas. To determine whether NDM could predict the atrophy pattern in ALS, the accumulation of pathology modelled by NDM was correlated against atrophy measured using MRI. The cross-sectional analyses revealed that the network diffusion seeded from the inferior frontal gyrus (pars triangularis and pars orbitalis) significantly predicts the atrophy pattern in ALS compared to controls. Whereas, atrophy over time with-in the ALS group was best predicted by seeding the network diffusion process from the inferior temporal gyrus at 6-month and caudal middle frontal gyrus at 12-month. Our findings suggest the involvement of extra-motor regions in seeding the spread of pathology in ALS. Importantly, NDM was able to recapitulate the dynamics of pathological progression in ALS. Understanding the spatial shifts in the seeds of degeneration over time can potentially inform further research in the design of disease modifying therapeutic interventions in ALS.

## Introduction

Amyotrophic Lateral sclerosis (ALS) is a progressive and fatal neurogenerative disorder, which involves the degeneration of both upper and lower motor neurons (Kiernan et al., 2011). Although ALS is primarily considered a disease affecting the motor system, extra-motor cerebral involvement in ALS is consistently reported (Foerster et al., 2013). The classical neuropathological features in ALS include the degeneration of the corticospinal tract (CST) and the grey matter in the spinal cord, brain stem and the motor cortex (Wang et al., 2011). The causes of the pathogenesis are largely unknown despite a range of studies indicating that neurodegenerative mechanisms include a complex interplay of excitotoxic stimulation, oxidative stress, neuroinflammation and the dysfunction of critical proteins as well as a wide range of genetic factors (Bonafede and Mariotti, 2017; Carri et al., 2003; Wang et al., 2011).

Neuropathological studies have shown the spreading of phosphorylated 43-kDa transactive response DNA-binding protein (pTDP-43) pathology in the central nervous system in ALS (Jo et al., 2020; Riku, 2020). Although the biomolecular mechanisms of trans-neuronal pTDP3 spread remain to be understood, it has been posited that the spread of abnormal proteins are primarily induced and disseminated from cortical neurons, via axonal connections, through synaptic contacts to the spinal cord and other brain regions (Braak et al., 2013). A key disease characteristic in ALS is the propagation of pTDP-43 pathology in regions far apart from each other (Braak et al., 2017). One hypothesis is that the corticospinal tract enables the propagation of pTDP-43 aggregates from the Betz cells in the agranular motor cortex to the distant (alpha) motor neurons in the spinal cord (Braak et al., 2017). These studies identify the role that axonal connections may play in pTDP3 spread, suggesting the hypothesis that the canonical organisation of axonal connections (human brain connectome) may determine the neuropathological changes that occur in ALS.

The spread of critical proteins in neurodegenerative diseases share similar pathological features. Besides ALS, the accumulation TDP-43 aggregates has been frequently observed in other neurodegenerative diseases including Frontotemporal Dementia (FTD) and Alzheimer’s disease (see a review paper by Jo et al., 2020) (Jo et al., 2020). In ALS, TDP-43 pathology is observed in more than 90% of the population (Jo et al., 2020). It has been proposed that the abnormal proteins in neurodegenerative diseases propagates in a “prion-like” manner (Jaunmuktane and Brandner, 2020). However, the key regions involved in the pathological propagation (often referred as the seed regions) are different in each disease. It has been identified that the normal connectivity profiles of these regions resemble the disease associated pattern of neural degeneration (Zhou et al., 2012).

Computational models have been developed to characterise how the organisation of the human connectome may facilitate the spread and accumulation of abnormal proteins in the brain. Raj et al., (2012) first demonstrated that the microscopic consequences of the pathological spread throughout the human brain can be modelled using a computational network diffusion model (NDM) (Raj et al., 2012). NDM has been extensively used to model the distribution of group-level pathology in a range of neurological diseases including Huntington’s disease (Poudel et al., 2019), dementia (Raj et al., 2012), Alzheimer’s disease (Raj et al., 2015), and Parkinson’s disease (Pandya et al., 2019). Poudel et. al., (2020) developed a personalised NDM to map subject-specific seeds of degeneration in traumatic brain injury (Poudel et al., 2020).

Two previous studies have used a similar computational approach to study network spread in ALS. Meier et al. (2020) showed that a computational random walker model can mimic progressive network degeneration in ALS (Meier et al., 2020). Their findings support the hypothesis that ALS impairment commences in the motor cortex and spreads along the white matter tracts to other brain regions. A NDM study by Sneha et al., (2021) showed that the basal ganglia, thalamus and insula, rather than the motor cortex, are likely to be involved in the pathological spread in ALS (Pandya et al., 2021). However, there is a lack of understanding of how the seeds of degeneration progress in ALS within a short timeframe (i.e., 6 to 12 months). This information is necessary for designing therapeutic interventions targeting the dissemination of ALS pathology.

In this study, we investigated whether NDM can determine the severity (compared to controls) and progression (with-in group changes over 12-months) of neurodegeneration in limb-onset ALS. We implemented NDM as per previously published framework (Poudel et al., 2019; Raj et al., 2012). We hypothesised that network diffusion via the human brain connectome is predictive of the severity and progression of the atrophy pattern observed in limb-onset ALS.

## Materials and Methods

### Participants

Ethics approval for this study was obtained from Calvary Health Care Bethlehem and Monash University Research Ethics Committees (Project Number: CF14/3968- 2014002057). Fourteen participants (n=14) with clinically possible, probable or definite limb-onset ALS and twelve (n=12) age matched control subjects underwent MRI scans. Twelve patients with ALS participated in follow-up MRI scans that were acquired six months after the initial scans. Nine out of twelve patients, who participated in the six month follow-up scan, also participated in a twelve-month follow-up MRI scan (Table 1). All the participants recruited in the study were over 18 years of age and able to give informed consent. Control subjects recruited for the study were screened for neurologic and psychiatric diseases. We were unable to collect the MRI data from all the individuals with ALS at follow-up scans due to their difficulty to lie flat inside the scanner.

**Table 1.** Participant demographics. Disease duration based on patient report of symptom onset to time of baseline scan.

|  | ALS | Controls |
| --- | --- | --- |
| Number of participants at baseline | 14 | 12 |
| Number of participants at six-months follow-up | 12 |  |
| Number of participants at twelve-months follow-up | 9 |  |
| Age (years) | 54.7 ± 13.0 | 51.4 ± 11.5 |
| Disease duration | 3.2± 3.8 |  |

### MRI data collection

MRI (T1-weighted) data for baseline and follow-up scans were acquired using a 3-Tesla Skyra MRI (Siemens, Erlangen, Germany) at Monash Biomedical Imaging, Melbourne, Australia. The T1-weighted magnetization prepared rapid gradient echo (MPRAGE) images were acquired using the following parameters: acquisition time = 5:20 minutes, repetition time = 2300ms, echo time = 2.07ms, flip angle = 9°, field-of-view = 256mm, voxel size = 1 × 1 × 1 mm^3^, 192 slices per volume.

### Structural connectome

The structural connectome data were obtained from the IIT Human Brain Atlas (V.5.0) (xhttps://www.nitrc.org/projects/iit/) and generated using artefact-free MRI data from 72 subjects (Qi and Arfanakis, 2021).

### Structural MRI data processing

Structural MRI (T1W) data for each subject were processed using the neuroimaging analysis tool, FreeSurfer version 6.0.0 (Fischl, 2012; Reuter et al., 2012). FreeSurfer was used to parcellate and segment the T1W data into 82 distinct cortical and sub-cortical brain regions based on the Deskian-Killiany atlas. We used the FreeSurfer longitudinal processing scheme, which is designed to process longitudinal MRI data i.e., data of the same individual at different time points. The neuroanatomical labels, after the cortical parcellation and sub-cortical segmentation, were visually inspected for accuracy in all ALS and control subjects.

### Network Diffusion Model

The NDM implementation was based on the previous work (Poudel et al., 2019; Raj et al., 2012). A brain network can be represented as a connectivity matrix *C* = {*c*_*i,j*_} where *c*_*i,j*_, represent the connection strength of white matter fibre pathways between *i*^*th*^ and *j*^*th*^ grey matter regions. In our study, the grey matter regions were obtained from the parcellation of the MRI T1-weighted images and the strength of fibre pathways between the nodes *c*_*i,j*_ were obtained from fibre tractography from the IIT Human Brain Atlas. The strength of fibre pathways is the number of fibre (streamlines) connected between nodes.

According to the Network Diffusion Model, the diffusion of pathological proteins through the white matter fibres connecting the grey matter brain regions at ***t*** time can be modelled as:

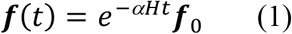

where *H* is the symmetric normalized Graph Laplacian. The time *t* is the model diffusion time, which has an arbitrary unit (a.u.). The vector ***f***_0_ represents the initial diffusion (i.e., at time zero). The term −*αHt* is known as the diffusion kernel. The graph Laplacian *H* can be represented as the equation below:

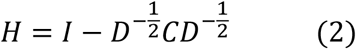

where *D* is a diagonal matrix, which contains the degree of each node. The degree of each node is defined as the sum of weighted connections emanating from that node. *I* is the identity matrix.

The computation of equation (1) can be obtained via eigenvalue decomposition of the Graph Laplacian *H* = *UΛ U*^*T*^, where *U* = [***u***_1_ … … ***u***_*N*_]. The computational solution of equation 1 is:

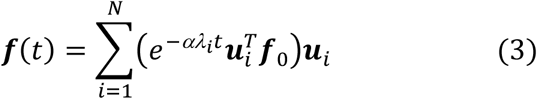

where *λ*_*i*_ and ***u***_*i*_ are the eigenvalues and eigenmodes of the of the Graph Laplacian *H*, and 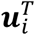 is the transpose of eigenmodes.

### Repeated seeding of network diffusion

Based on the repeated seeding approach by Poudel et al. (2019) we implemented NDM (equation 3) to identify the epicentre of the disease. For each brain region, the network diffusion model was used to estimate the diffusion at all other regions at time *t* = 0 to 50. Network diffusion was initiated from bilateral seeds, such that 41 different initial conditions were used. This process generated vectors with 80 elements at each time point. The NDM vector at all time points, ***f***(*t*), were correlated against measured atrophy (t-value) using Pearson’s correlation, resulting in 41 × 51 correlation-time matrices. It is noted that the correlation was performed excluding the seed regions, in order to ensure that the correlation was not driven by the atrophy in the seed region (Poudel et al., 2019). The maximum correlation value for each seed region was used to estimate the likelihood of that region being involved in the spread of the pathology in ALS.

### Estimation of measured atrophy (t-statistics)

To measure the atrophy in the parcellated brain regions, the mean and standard deviation (SD) of the volume were estimated. Regional volumes were normalized using total intracranial volume to correct for head size (Whitwell et al., 2001). For each region, t-statistics of the volumetric difference between the ALS patient group and the control group were estimated. The t-statistics were measured between control subjects and ALS patients at baseline, ALS patients at baseline and at six-month follow-up, and ALS patients at six-month follow-up and at one-year follow-up. The two sample t-test was used when computing the t-statistics between the control subjects and ALS patients at baseline. Paired sample t-tests were used to compute the t-statistics between the ALS patients at baseline and at six-month follow-up, and between six-month follow-up and one-year follow-up. The p value significance level for t-tests was 0.05. Uncorrected p-values are reported as NDM modelling uses effect-sizes (t-values) to measures the strength of atrophy.

### Correlation with NDM

The measured atrophy (t-scores) were correlated with the predicted atrophy (i.e., NDM vector ***f***(*t*)) using Pearson’s correlation, as described in the “ Repeated seeding of network diffusion” section. Correlation values with p values (family-wise corrected based on brain regions) less than 0.05 were considered to be significant.

## Results

### Measured atrophy pattern in ALS

#### At baseline (cross-sectional)

In ALS patients at baseline compared to control subjects, volume loss was observed in brain regions including the right pars triangularis (t = 2.64, p = 0.01), the right superior frontal gyrus (t = 2.19, p = 0.04) and the left lateral orbitofrontal gyrus (t = 2.12, p = 0.04).

#### At six-month and twelve-month follow-up (longitudinal)

In ALS patients at six-months compared to baseline, the five most prominent brain regions showing volumetric loss were the left lateral occipital gyrus (t = 3.56, p = 0.005), right inferior temporal gyrus (t = 3.1, p = 0.01), left inferior temporal gyrus (t = 2.97, p = 0.01), right middle temporal gyrus (t = 2.93, p = 0.01) and left fusiform gyrus (t = 2.88, p = 0.02).

In ALS patients at twelve-months compared to six-months, volume loss was observed in the brain regions including right bankssts (cortical areas around superior temporal sulcus) (t = 3.5, p = 0.008), left pars opercularis (t = 2.89, p = 0.02), left putamen (t = 2.86, p = 0.02), right precentral gyrus (t = 2.39, p = 0.04) and left caudal anterior cingulate (t = 2.34, p = 0.05). For visualization, the measured atrophy pattern (t-scores) for both cross-sectional and longitudinal analyses is displayed in Figure 1.

**Figure 1.**
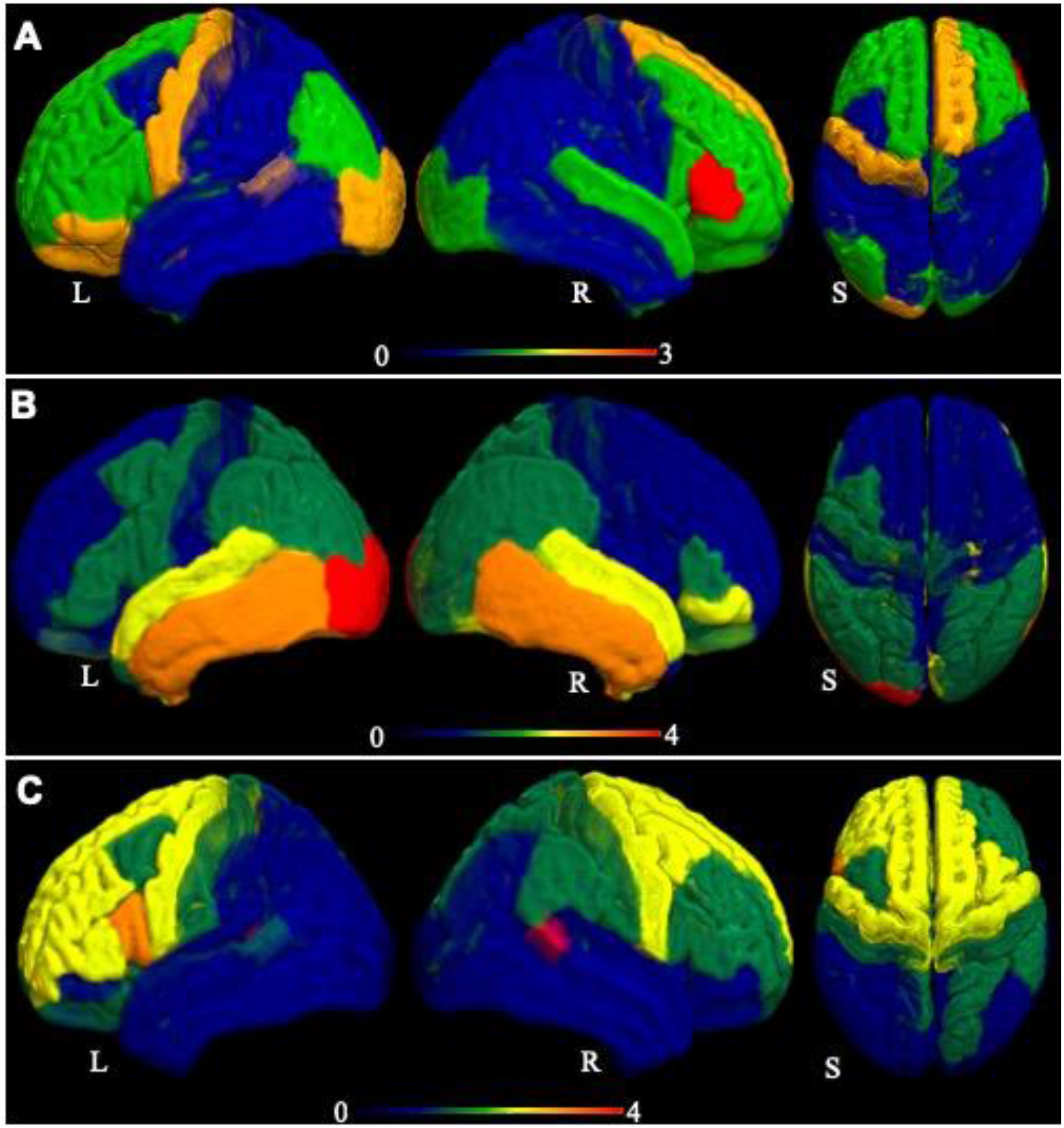
Visual representation of the measured atrophy in ALS patients. (A) The measured atrophy at baseline compared to controls, (B) at six months follow-up compared to baseline (B), and (C) at twelve-months follow-up compared to six-month follow-up. The displayed atrophy pattern is based on the t-value of the difference in grey-matter volume in the 82 parcels from the Desikan-Killiany atlas. The darker red colour in the colour range represents the region of greatest atrophy. L, R and S represent left, right, and superior view.

### Network diffusion model of atrophy pattern in ALS

#### At baseline

The maximum correlation between the measured atrophy at baseline (ALS versus controls) and predicted atrophy was observed when the diffusion was initiated from two brain regions, namely the pars triangularis (r = 0.41, p = 0.02, corrected) and the pars orbitalis (r = 0.39, p = 0.02, corrected) (Table 2). Figure 2 (A) shows the linear association between the measured and predicted atrophy, whilst Figure 2 (B) provides a visual representation of the maximum correlation obtained at each node, for baseline.

**Table 2:**
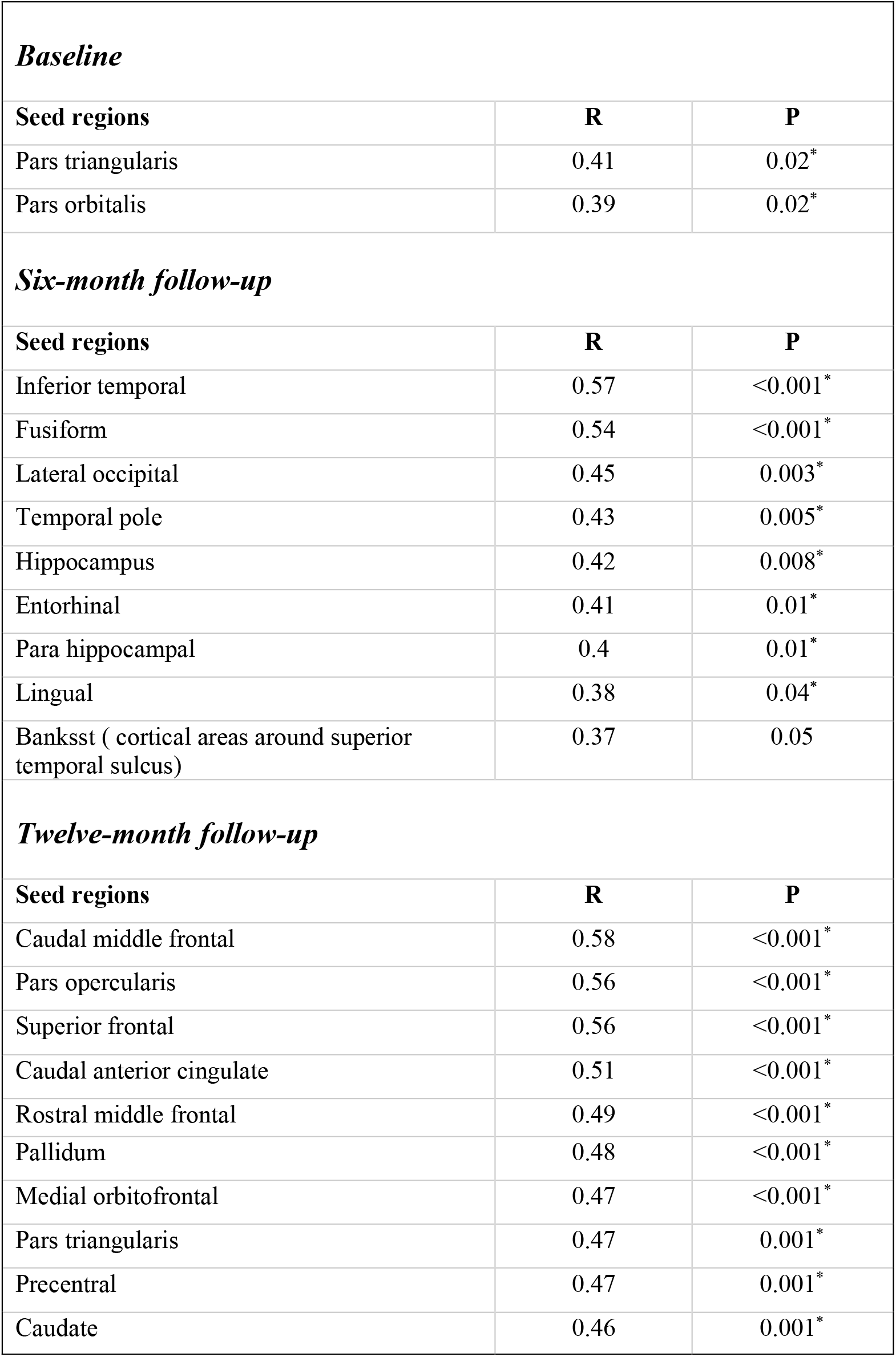
Seed regions identified by NDM at baseline (atrophy measured at baseline compared to controls), at six-month follow-up (atrophy measured at six-month follow-up compared to the baseline) and at twelve-month follow-up (atrophy measured at twelve-month follow-up compared to the six-month follow-up). R is Pearson’s correlation. p values are family-wise error (FWE) corrected). The symbol * denotes significant p value.

|  |  |  |
| --- | --- | --- |
| ***Baseline*** |  |  |
| **Seed regions** | **R** | **P** |
| Pars triangularis | 0.41 | 0.02* |
| Pars orbitalis | 0.39 | 0.02* |
| ***Six-month follow-up*** |  |  |
| **Seed regions** | **R** | **P** |
| Inferior temporal | 0.57 | <0.001* |
| Fusiform | 0.54 | <0.001* |
| Lateral occipital | 0.45 | 0.003* |
| Temporal pole | 0.43 | 0.005* |
| Hippocampus | 0.42 | 0.008* |
| Entorhinal | 0.41 | 0.01* |
| Para hippocampal | 0.4 | 0.01* |
| Lingual | 0.38 | 0.04* |
| Banksst ( cortical areas around superior temporal sulcus) | 0.37 | 0.05 |
| ***Twelve-month follow-up*** |  |  |
| **Seed regions** | **R** | **P** |
| Caudal middle frontal | 0.58 | <0.001* |
| Pars opercularis | 0.56 | <0.001* |
| Superior frontal | 0.56 | <0.001* |
| Caudal anterior cingulate | 0.51 | <0.001* |
| Rostral middle frontal | 0.49 | <0.001* |
| Pallidum | 0.48 | <0.001* |
| Medial orbitofrontal | 0.47 | <0.001* |
| Pars triangularis | 0.47 | 0.001* |
| Precentral | 0.47 | 0.001* |
| Caudate | 0.46 | 0.001* |

**Figure 2.**
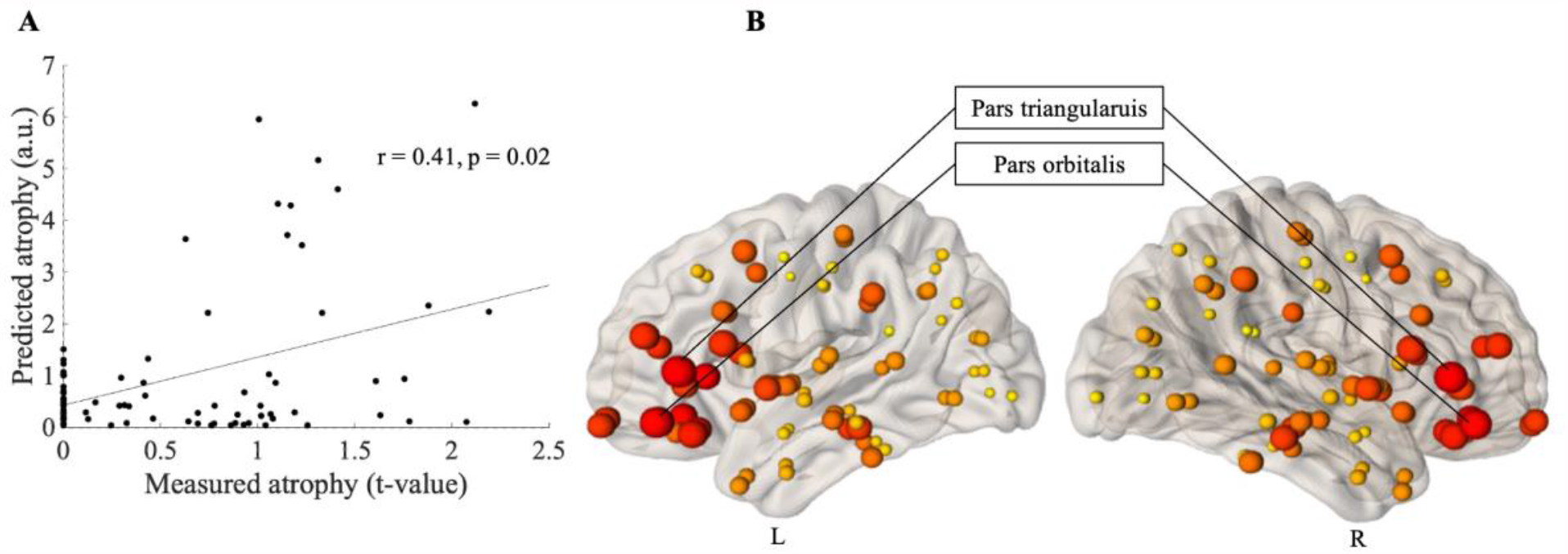
Prediction of atrophy pattern at baseline. (A) Scatter plot of the correlation between predicated and measured atrophy, when measured atrophy was estimated between ALS patients at baseline and control subjects. Pearson’s correlation coefficient (r) and the significance of the correlation (p) are shown, p is family-wise error (FWE) corrected. (B) Visual representation of maximum correlation obtained for each node, when measured atrophy was estimated between ALS patients at baseline and control subjects. The size of the ball corresponds to the maximum correlation between predicted and measured atrophy. The colour of the balls ranges from light yellow (for smallest) to dark red (for biggest). L is left and R is Right.

#### At six-month follow-up

The maximum correlation between the progression of atrophy over 6-months within the ALS. Group and NDM predicted atrophy was observed when the diffusion was initiated from brain regions including inferior temporal gyrus (r = 0.57, p < 0.001, corrected), fusiform gyrus (r= 0.54, p < 0.001, corrected) and the lateral occipital and temporal pole (r= 0.44, p= 0.003, corrected) (Table 2). Figure 3A shows the linear association between the measured and predicted atrophy, whilst Figure 3B provides a visual representation of the maximum correlation obtained at each node, for six-month follow-up.

**Figure 3.**
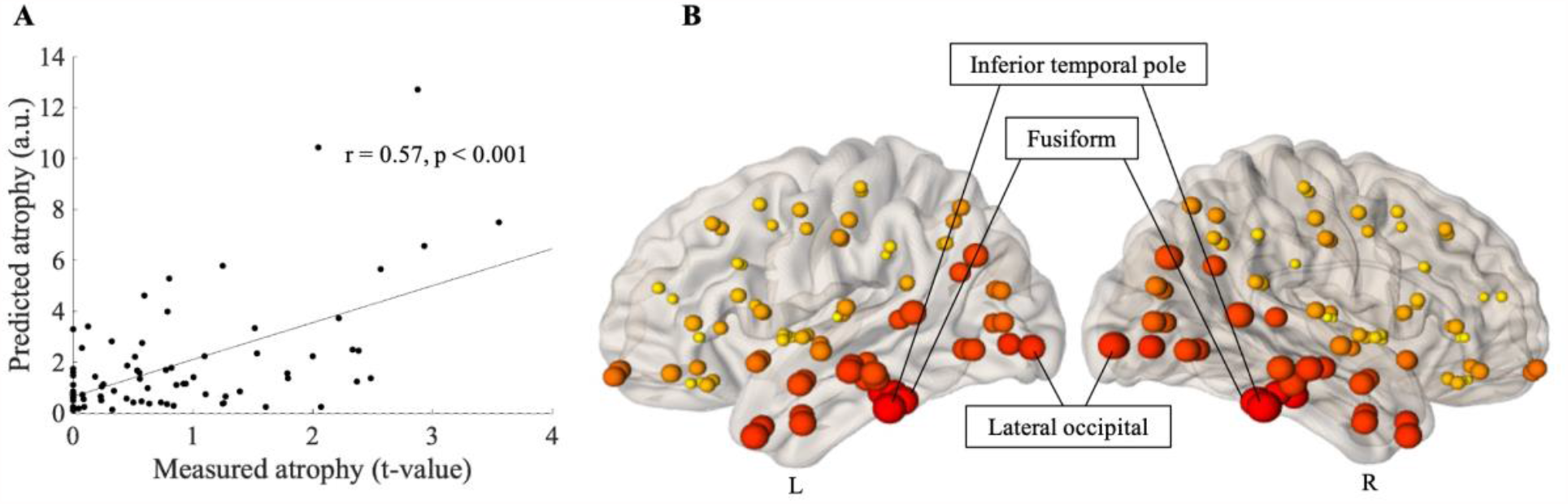
Prediction of atrophy pattern at six-month follow-up. (A) Scatter plot of the correlation between predicated and measured atrophy, when measured atrophy was estimated between ALS patients at six-month follow-up and ALS patients at baseline. Pearson’s correlation coefficient (r) and the significance of the correlation (p) are shown, p is family-wise error (FWE) corrected. (B) Visual representation of maximum correlation obtained for each node, when measured atrophy was estimated between ALS patients at six-month follow-up and ALS patients at baseline. The size of the ball corresponds to the maximum correlation between predicted and measured atrophy. The colour of the balls ranges from light yellow (for smallest) to dark red (for biggest biggest). L is left and R is Right.

#### At twelve-month follow-up

When the measured atrophy was estimated between the ALS patients at six-month follow-up and at twelve-month follow-up, the maximum correlation between the measured and predicted atrophy was observed when the diffusion was initiated from brain regions including the caudal middle frontal (r = 0.58, p < 0.001, corrected), pars opercularis (r= 0.56, p < 0.001, corrected), superior frontal gyrus (r = 0.56, p < 0.001, corrected) and caudal anterior cingulate (r = 0.51, p < 0.001, corrected). The top ten seed regions identified by NDM are shown in Table 2.

Figure 4A shows the linear association between the measured and predicted atrophy, whilst Figure 4B provides a visual representation of the maximum correlation obtained at each node, for twelve-month follow.

**Figure 4.**
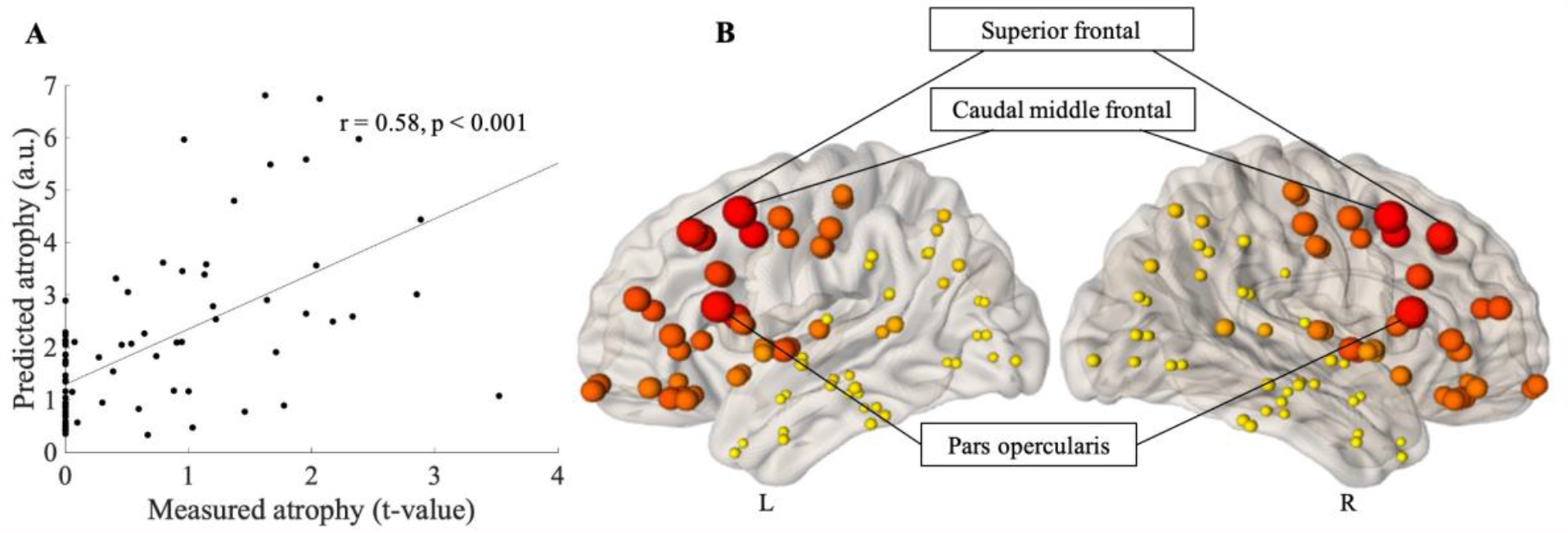
Prediction of atrophy pattern at twelve-month follow-up. (A) Scatter plot of the correlation between predicated and measured atrophy, when measured atrophy was estimated between ALS patients at twelve-month follow-up and ALS patients at six-month follow-up. Pearson’s correlation coefficient (r) and the significance of the correlation (p) are shown, p is family-wise error (FWE) corrected. (B) Visual representation of maximum correlation obtained for each node, when measured atrophy was estimated between ALS patients at twelve-month follow-up and ALS patients at six-month follow-up. The size of the ball corresponds to the maximum correlation between predicted and measured atrophy. The colour of the balls ranges from light yellow (for smallest) to dark red (for biggest biggest). L is left and R is Right.

## Discussion

In this study, we implemented a computational model of network diffusion to determine whether the severity and progression of neurodegeneration in limb-onset ALS can be modelled as a diffusion process. Our results can be summarized into three main findings. First, a volumetric analysis of the longitudinal MRI data from ALS patients revealed progressive neurodegeneration in extra-motor brain regions. Second, the NDM applied on a canonical structural brain connectome was able to recapitulate the cross-sectional and longitudinal pattern of spatial neurodegeneration in limb-onset ALS. Third, NDM revealed spatial shifts in the seeds of degeneration over time. These findings demonstrate the utility of connectome based model of network diffusion in predicting the evolution of neurodegeneration in limb-onset ALS.

A cross-sectional comparison of the volumes within 82 brain regions between ALS patients at baseline and a group of healthy controls revealed degeneration in the right pars triangularis and superior frontal gyrus. This was, perhaps, not surprising given that ALS is a neurodegenerative disorder. Earlier structural MRI studies have consistently shown grey and white matter alterations in ALS (Bhattarai et al., 2020; Bhattarai et al., 2021; Chiò et al., 2014; Foerster et al., 2013). Voxel based morphometry studies have shown significantly reduced grey matter density in motor and premotor brain regions (Agosta et al., 2007; Turner et al., 2007). Other groups have reported atrophy in the frontotemporal regions (Mezzapesa et al., 2007). One study reported no significant grey matter atrophy in ALS patients compared to controls (Abrahams et al., 2005).

More importantly, we observed a significant loss of brain volume within the ALS group within 6-month, which further progressed in 12-month. At 6-month, degeneration was observed in the left lateral occipital gyrus, and bilaterally in inferior temporal brain regions, which further progressed to the right bankssts (cortical areas around superior temporal sulcus), left pars opercularis, and left putamen at twelve-month follow-up. While only a limited number of studies have characterised ALS degeneration within a short time frame, most of these studies have shown a complex pattern of neurodegeneration involving multiple brain systems. These findings include progressive volumetric loss in the frontal grey matter (Agosta et al., 2009), cortical thinning in the right pars-triangularis and left lateral orbital frontal region (Verstraete et al., 2012), grey matter volumetric loss in the left insula and right temporo-polar cortex, and white matter volumetric loss in the inferior frontal gyrus at baseline with pronounced bilateral frontal and temporal involvement at six-month follow up (Senda et al., 2011). Other neuroimaging studies have consistently shown the involvement of frontal and temporal regions in the disease process in ALS (see a review paper by Foerster et al., 2013) (Foerster et al., 2013). Taken together, these findings suggest vulnerability of both extra-motor and motor brain regions in ALS.

To understand the macroscopic processes driving the spatial pattern and severity of degeneration in ALS, we used a computational Network Diffusion Model as per the previously published framework (Poudel et al., 2019; Raj et al., 2012). We found that NDM was only moderately able to capture the spatial pattern of degeneration in ALS, particularly when compared against controls. The NDM was able to better recapitulate the longitudinal progression of degeneration within the ALS group. Interestingly, we observed a spatial shift in the most significant seed regions as the disease progressed. When degeneration maps of ALS patients compared to controls was used, the NDM identified frontal brain regions including the pars triangularis and pars orbitalis to be the most significant seed regions for the disease spread. At six-month follow-up, when the atrophy was estimated relative to baseline, temporal brain regions including the inferior temporal and fusiform were the most significant seed regions. At twelve-month follow-up, as the disease progressed further, the NDM identified the caudal middle frontal, pars opercularis and the superior frontal gyrus as the most significant seed regions. Such spatial shift in disease seeds is consistent with rapid progression of ALS and multi-network degeneration hypothesis in ALS.

It is important to note that both cross-sectional and longitudinal analysis reveal the frontal and temporal brain regions as most likely seeds to be involved in trans-neural disease progression in ALS. Perhaps surprisingly, we did not observe the motor cortex as a region from which the disease spread was initiated. However, the average duration of disease in our cohort was approximately three years and we may be observing disease progression rather than disease onset. An earlier cross sectional NDM study in ALS patients showed that the basal ganglia, thalamus and insula, rather than the motor cortex are likely to be involved in the pathological spread in ALS (Pandya et al., 2021). Conversely, a recent study based on a computational random walker model supports the conventional hypothesis that ALS impairment begins in the motor cortex and spreads to other brain regions in a spatiotemporal manner (Meier et al., 2020).

Post-mortem findings suggest that pTDP-43in ALS may disseminate in a sequential manner to disparate brain regions via axonal pathways, which underpins the identification of four neuropathological stages in the disease process (Brettschneider et al., 2013). The initial stage (Stage 1) is characterized by lesions developed in the agranular motor cortex, brainstem and motor nuclei. In Stage 2, the affected regions are located in the prefrontal neocortex, the brain stem (reticular formation), the pre-cerebellar nuclei and the red nucleus. In Stage 3, the affected regions are located in the prefrontal regions and then the postcentral neocortex and striatum. The final stage (Stage 4) has affected regions located in the anteromedial portions of the temporal lobe and the hippocampus. Although our longitudinal findings showed the involvement of frontal and temporal regions, they did not precisely overlap with the progressive pattern at different stages as described in the ALS staging scheme. The heterogeneity in the disease stage may be one of the potential reasons that our findings differed from the ALS staging scheme. Using a subject specific Network Diffusion Model, as shown in an recent traumatic brain injury study (Poudel et al., 2020), may be a useful approach for future ALS studies to precisely estimate the disease progression in an individual patient level.

Although TDP-43 pathology is observed in more than 90% of ALS population, there are discrepancies in findings related to the spread of TDP-43 inclusions in the central nervous system in ALS. Post-mortem findings suggest that TDP-43 aggregates spreads in the brain via neural projections (Braak et al., 2013; Brettschneider et al., 2013), whereas experimental findings have identified the cell-to cell propagation of aggregated TDP-43 in ALS (Jo et al., 2020; Riku, 2020). It remains to be clarified whether the propagative nature of TDP-43 aggregates plays a role in anatomical spreading of TDP-43 pathology in the central nervous system in ALS. The spreading of TDP-43 varies among the ALS patients, which indicates the need to examine different potential confounding host factors i.e. genes (Riku, 2020). This variability in the pathological spread could be one of the reasons that the epicentre of the disease identified in our study and other computational studies are different.

There are a number of limitations that need to be taken into consideration when interpreting the results. The sample size in our study is relatively small. Although we sought to minimize the heterogeneity using patients with limb-onset ALS, the cohort consisted of patients with different stages of disease severity. We were not able to recruit all of the patients scanned at baseline for follow-up scans, which further reduced the sample size for longitudinal analyses. The genetic factors and the patient’s medication history were not taken into consideration in the analysis. NDM is a first order linear model which does not take into account non-linear changes in the structural connectivity as the disease progresses. Furthermore, the NDM uses a healthy connectome from a healthy control group based on the assumption that structural network serves as a pathway for pathological spread, and remains unimpaired during the disease process (Torok et al., 2018). An interesting possibility would be for future NDM studies to incorporate connectome models that become impaired during the disease staging.

In summary, our findings suggest that the network diffusion model using the structural connectome may be a suitable tool for predicting severity and progression of degeneration in ALS. We showed that NDM can identify the brain networks that are responsible for pathological spread in ALS. In this study consisting of individuals with limb-onset ALS, extra-motor, frontal and temporal regions were more likely to be involved as the seed regions for network spread in ALS. Further studies with a focus on implementation of the NDM to investigate the degenerative pattern in individual patients may assist in addressing some of the issues identified in our study.

## CRediT author statement

**Anjan Bhattarai:** Conceptualization, Methodology, Validation, Software, Formal analysis, Investigation, Data curation, Writing – Original Draft, Writing – Review & Editing, Visualization. **Zhaolin Chen:** Methodology, Validation, Writing – Review & Editing, Supervision. **Phyllis Chua:** Investigation, Resources, Writing – Review & Editing, Supervision, Funding acquisition. **Paul Talman:** Investigation, Resources, Writing – Review & Editing, Supervision, Funding acquisition. **Susan Mathers:** Investigation, Resources, Writing – Review & Editing, Funding acquisition. **Caron Chapman:** Resources, Writing – Review & Editing. **James Howe:** Resources, Writing – Review & Editing. **Sarah Lee:** Resources, Writing – Review & Editing. **Yenni Lie:** Resources, Writing – Review & Editing. **Govinda R Poudel:** Conceptualization, Methodology, Software, Validation, Resources, Writing – Review & Editing, Visualization. **Gary F Egan:** Conceptualization, Resources, Writing – Review & Editing, Supervision.

## Acknowledgement

We would like to thank all the volunteers who participated in this study. We are also grateful to the radiographers at Monash Biomedical Imaging for their assistance with MRI data collection and our nurse research assistants Ruth Krasniqi and Anna Smith. Funding for this project was obtained through the Monash University Strategic Grant Scheme. AB was supported by The Australian Rotary Health / Rotary Club of Sandy Bay PhD Scholarship in Motor Neuron Disease. We declare no conflict of interest.

